# Genomic, physiological, and transcriptomic mechanisms of pH stress adaptation and response in *Methylocystis sumavensis*, a novel peatland methanotroph

**DOI:** 10.1101/2025.08.29.672306

**Authors:** Justus Amuche Nweze, Magdalena Wutkowska, Ghadir Ibrahim, Anne Daebeler

## Abstract

Methanotrophic bacteria in peatlands mitigate emissions of methane (CH_4_), a potent greenhouse gas, yet the mechanisms enabling them to remain active under the acidic conditions typical of many peatlands remain poorly understood. Using enrichment cultivation and single-cell sorting, we isolated a novel peatland methanotroph from Czech soil, ‘*Methylocystis sumavensis*’. This species is moderately acidotolerant and active across broad pH and temperature ranges, with growth optima at pH 6.8 and 24–37 °C. The genome of ‘*M. sumavensis*’ encodes two particulate methane monooxygenase isozymes, a clade I nitrous oxide reductase, multiple terminal oxidases, and two [NiFe]-hydrogenases, including a complete complex of the previously uncharacterised membrane-bound hydrogenase group 4f, indicating substantial metabolic versatility. To link genomic potential with physiological function, we compared transcriptomes under acidic (pH 5.0) and alkaline (pH 9.0) conditions relative to the optimum pH (6.8). Under both stresses, ‘*M. sumavensis*’ increased transcription of genes involved in membrane remodelling, ion transport across both membranes, and stress response and repair, while reducing transcription of genes associated with methane oxidation and cell division. Alkaline stress additionally suppressed growth through reduced transcription of the carbon-assimilating Serine cycle. In contrast, acidic stress triggered a coordinated response requiring greater energetic investment. Among the most strongly upregulated genes were those encoding two formate dehydrogenases, the branched-chain alpha-keto-acid dehydrogenase complex, and all seven group 4f [NiFe]-hydrogenase-related genes, suggesting enhanced respiratory redox balancing under elevated external proton concentrations. These results reveal key mechanisms of pH-stress adaptation and highlight metabolic plasticity as a major determinant of methanotroph resilience in ecosystems.

## Introduction

Methane (CH_4_) is a potent greenhouse gas with a global warming potential 81.2 times greater than that of carbon dioxide (CO_2_) on a 20-year timescale [1]. Since pre-industrial times, atmospheric CH_4_ concentrations have almost tripled, now exceeding 1,934 parts per billion [2], driven by both anthropogenic and natural emissions [3]. Wetlands, including peatlands, are the largest natural biogenic source of CH_4_ and have been considered the main source of the recent CH_4_ growth rate anomaly [4, 5]. Nevertheless, peatlands are a net carbon (C) sink [6]. Despite covering only 3% of Earth’s terrestrial surface [7], peatlands contain about 25% of the global soil C stock [8], which is equivalent to twice the amount in the world’s forests [9]. Alarmingly, anthropogenic pressure on peatlands is increasing [10] and experts predict peatlands to become a net C source within this century [11].

Peatlands support methanotrophic communities capable of mitigating CH_4_ emissions by oxidising it before it escapes into the atmosphere [12]. Methanotrophs are bacteria that can use CH_4_ as their sole C and energy source [13], and peatlands typically harbour abundant and compositionally distinct methanotroph communities [14–16]. Among known aerobic methanotrophs, the *Proteobacteria*, particularly the alphaproteobacterial family *Methylocystaceae*, typically dominate in acidic peatlands [12, 17, 18]. Members of this family, including the genus *Methylocystis*, are characterised by facultative methanotrophy and mixotrophy [19–22]. Despite their ecological importance, these organisms remain notoriously fastidious and only a few peatland *Methylocystis* sp. have been brought to pure culture [23, 24], limiting our understanding of the specific molecular adaptations that facilitate their dominance within methanotrophic communities in acidic ecosystems and predictions on how CH_4_-cycling communities will respond to environmental change [25]. Acidic environments impose distinct bioenergetic constraints, which lower the maximum energy conservation yield of resident microorganisms [26, 27]. Moreover, a lowered intracellular pH caused by proton influx can lead to protein instability, reduced enzymatic activity, and structural damage of both DNA and RNA, making the maintenance of cytoplasmic pH homeostasis essential for cellular function [28]. However, since sustaining near-neutral intracellular pH against a steep transmembrane proton gradient requires additional energy, not only are specialised adaptations in membrane composition and ion transport are essential, but also flexible and efficient energy conservation mechanisms that meet the increased energy demand [27]. To date, the mechanisms enabling high methanotroph abundance and activity in acidic peatlands are not well understood [29]. Genomic and lipidomic studies of methanotrophs from other acidic systems have implicated antiporters, membrane potential-based proton exclusion, stress-responsive chaperones, and increased saturated phospholipid contents in acid stress responses [29, 30]; however, evidence that peat methanotrophs use these mechanisms is currently lacking.

Previously, we reported that *Methylocystis* spp. dominate the methanotrophic community in an acidic minerotrophic fen in the Šumava National Park (Czechia), and phylogenetic analyses suggested the presence of uncultured lineages [18]. Here, we describe the isolation and characterisation of a novel acid-tolerant *Methylocystis* species, ‘*Methylocystis sumavensis’*, from this site. Using a combination of single-cell sorting, whole-genome sequencing, metabolic reconstruction and comparative genomics, physiological assays across a pH gradient (5.0–9.0), and transcriptomic profiling under acidic (pH 5.0), neutral (pH 6.8), and alkaline (pH 9.0) conditions, we define its physiological limits of CH_4_ oxidation and link genomic potential with condition-specific transcriptional responses to acid and alkaline stress. Our findings expand the current understanding of methanotroph ecology in acidic peatlands and provide the first insights into methanotroph responses to environmental change and mechanisms of persistence in carbon-rich, acidic, and nutrient-poor peatland ecosystems.

## Materials and Methods

### Sampling, enrichment, and isolation

Peat samples were collected from the surface layer (1–15 cm) of a pristine minerotrophic fen in the Šumava National Park, Czechia. The site is naturally acidic, with a pH range of 3.8–4.2, and has been described in detail previously [31, 32]. Before preparing enrichment cultures, 150 g of unmixed peat soil from the top 15 cm was incubated statically at 24°C in a sterile butyl rubber-capped 500 ml glass bottle amended with 5% CH_4_ for one week. Afterwards, 0.2-0.5 g of peat were placed into wells of a 24well culture plate (Greiner) containing 1.5 ml of M2 medium [33]. The plate was placed into air-tight incubation boxes equipped with ports for gas sampling (Don Whitley Scientific), adjusted with 5% CH_4_, and incubated statically at 24°C for 2 weeks. Wells were then screened for methanotroph presence by PCR and subsequent gel electrophoresis using primers pmoC-374F/ pmoA-344R [34] with 2 µl enrichment culture and the following reaction conditions: initial denaturation at 96 °C for 5 min, followed by 35 cycles at 96 °C for 1 min, annealing at 55 °C for 1 min, and elongation at 72 °C for 2 min. The final elongation step was done for 10 min at 72 °C. Material from positive wells was used for serial dilution ranging from 10^-2^ to 10^-4^ in M2 and NMS [35] medium incubated in sterile 100 ml glass bottles capped with grey butyl rubber stoppers for two weeks as described above. CH_4_ concentrations in the headspace were monitored using gas chromatography (Agilent Technologies 6850 GC System; see details below). As enrichments in NMS medium showed higher activity, this medium was chosen for incubation from here on. After consumption of 70-90% of the provided CH_4_, 100 µl of enrichment culture was sampled and used for inoculation of 50% NMS agar plates either directly or diluted 100-fold. Plates were incubated in air-tight incubation boxes as described above. After 2.5 weeks, small transparent colonies were picked and streaked onto new plates three consecutive times before inoculating liquid medium. Incubation and monitoring of CH_4_ consumption was performed as before, but attempts to re-grow on agar plates failed. The enrichment culture was then preserved at -70°C in 10% DMSO before re-activation and single-cell sorting to isolate methanotrophs from the enrichment culture. Specifically, the enriched cells were randomly sorted into sterile 96-well tissue culture plates (Life Science) without the use of fluorescent dyes by using a fluorescence-activated cell sorting (FACS) instrument (FACSAria™ III, 70 μm nozzle, 70 psi) equipped with FACSDiva version 8 software (BD Biosciences). Cells were visualised in the forward and side scatter channels and small, non-cell particles as identified in sterile medium were excluded; otherwise, no gating was applied. The wells of the microtiter plates contained 200 μl of sterile NMS medium. The plates were then placed in air-tight incubation boxes equipped with ports for gas sampling (Don Whitley Scientific) and incubated at 24°C in the dark without agitation after the addition of ∼10% CH_4_ to the headspace. After 30 days, turbid wells were screened for taxonomic identity and purity via Sanger sequencing after colony PCR (16S rRNA gene amplification using primers 27-F [5-AGAGTTTGATCCTGGCTCAG-3] and 1492-R [5-GGTACCTTGTTACGACTT-3]) [36] and PCR product cleaning with the ExoSAP_IT_express kit (Applied Biosystems™) according to the manufacturer’s instructions. Cultures from wells that produced clean 16S rRNA gene sequences (only unambiguous base peaks in the Sanger chromatogram) that were affiliated to known methanotroph genera were transferred (1:1000) into 100 ml fresh, sterile NMS medium and incubated as described above. Culture purity was routinely verified by streaking on LB agar (MP Biomedicals) and incubation at 20°C for 10 days. Details of methods employed for scanning electron microscopy, DNA extraction and genome sequencing, and genomic data processing are given in the Supplementary Text.

### Phylogenomics, annotation, and gene-specific phylogenies

Public genomes of 38 cultured and 45 uncultured *Methylocystis* strains were downloaded from NCBI. The quality was assessed as described above, and only 40 high-quality genomes and MAGs (> 92% completeness, 9% contamination) were retained for downstream analyses together with the circular genome of our new isolate (Table S1). Single-copy core genes (SCGs, n = 71) were identified with *anvi-run-hmms* and extracted as amino acid sequences with *anvi-get-sequences-for-hmm-hits* in Anvi’o (v8.0) [37–39]. Protein sequences of the SCGs were aligned with MAFFT using the *--auto* function [40] and used to construct a maximum-likelihood phylogenomic tree using IQ-TREE v2 [41] with the LG+F+R3 substitution model selected by the implemented Modelfinder function [42] and 1,000 bootstrap replicates.

Genomes were annotated using DRAM (Distilled and Refined Annotation of Metabolism) [43]. Unannotated genes of interest, especially for multi-subunit proteins (such as nitrous oxide reductase, cytochrome c oxidases, and ATP synthases) were manually cross-validated using the annotation by Clusters of Orthologous Groups (COGs) database via BLASTP implemented in Anvi’o [44]. In addition, selected gene families were re-screened using HMM profiles derived from curated reference proteins where appropriate [45]. In cases where annotations remained ambiguous or incomplete, additional BLASTP searches were performed manually against the NCBI nr database to confirm gene identity. Final functional assignments were based on consensus across tools to ensure accuracy in metabolic reconstruction. Additional annotation efforts to assess the specificity of the H⁺/Na⁺-translocating ATP synthase and the methodology of gene-specific phylogenies are described in the Supplementary Text.

### Optimal conditions for CH_4_ oxidation

To assess the effects of temperature on the activity of the newly isolated *Methylocystis* species, cells were harvested in exponential phase by centrifugation at 10,000 x g for 10 min, washed twice, and resuspended in 80 ml of fresh medium. Aliquots of culture (4 ml) were transferred to sterile glass tubes capped with butyl rubber stoppers. After replacing ∼10% of the headspace volume with 99% pure CH_4_, the tubes were incubated in triplicate at 4°C, 15°C, 24°C, 30°C, 37°C, and 42°C, shaking at 100 rpm. CH_4_ concentrations in the headspace were measured initially and at regular intervals (at least five times between 2 and 52 h after start of the incubation). CH_4_ oxidation rates were calculated assuming either zero-order or first-order kinetics, depending on the best linear fit to the time-resolved concentration data according to the Akaike Information Criterion.

For determination of the pH range and optimum for activity of the newly cultured methanotroph, pre-grown cultures were harvested, washed, and resuspended as described above into 20 ml of fresh NMS medium adjusted to pH 4.0, 5.0, 6.8, 8.0, or 9.0 in triplicate. pH was controlled using a low-phosphate buffer system (KH_2_PO₄ and K_2_HPO₄; final total phosphate concentration 4 mM), with fine adjustment using H_3_PO_4_ or KOH as required. For pH 9.0, KOH was used for final adjustment following phosphate addition. A detailed description of buffered NMS preparation is provided in the Supplementary Methods. The headspace of the sealed bottles was replaced with ∼10% (v/v) CH_4_ and incubated at 24°C with shaking at 100 rpm. CH_4_ concentrations in the incubation headspace were monitored repeatedly over time, with a minimum of five measurements within the first 30 h for all pH treatments, except pH 4.0, where activity was followed for up to 137 h. Stability of pH was verified during the incubation using contactless optical pH sensor spots (ranges 4.0–6.0 and 7.0–9.0) and a FireSting-PRO optical meter (PyroScience) according to the manufacturer’s instructions. For biomass quantification (total protein), 200 µl of culture was collected from all replicates before and after incubation at each different pH and temperature condition and stored at −20 °C. After the incubation, cultures maintained at pH 5.0, 6.8, and 9.0 were harvested by centrifugation at 10,000 x g and 4°C, and immediately frozen at −70 °C for transcriptomic analyses. Importantly, cultures were still actively oxidising CH_4_ at the time of sampling.

The setup of the assay used to test molecular hydrogen production by ‘*M. sumavensis’* under acidic conditions is described in the Supplementary Text.

### CH_4_, carbon dioxide, and molecular hydrogen quantification

CH_4_ and CO2 concentrations were measured using a gas chromatograph (Agilent Technologies 6850 GC System) equipped with a flame ionisation detector and a ShinCarbon ST Micropacked column (2 m, 0.53 mm ID, Restek). CH_4_ concentrations (μM) were determined from gas measurements by applying the ideal gas law at standard conditions. The quantification of molecular hydrogen is described in the Supplementary Text.

### Total RNA extraction and sequencing

Total nucleic acids were extracted from cultures incubated at pH 5.0, 6.8, and 9.0 during active CH_4_ oxidation and prior to substrate depletion, following the protocol described in [46]. Extracts were treated with TURBO™ DNA-free DNase (Invitrogen) to remove residual genomic DNA, and total RNA was subsequently purified using the GeneJET RNA Purification Kit (Thermo Fisher Scientific). The absence of contaminating DNA was confirmed by the failure to amplify the 16S rRNA gene by PCR using primers 27-F and 1492-R as described in [36]. RNA concentration was quantified using the Qubit™ RNA HS Assay Kit on a Qubit™ 4 Fluorometer (Invitrogen). Purified RNA samples were sequenced by Novogene (Munich, Germany) using a prokaryotic strand-specific mRNA-seq workflow with rRNA removal. Libraries were sequenced on an Illumina NovaSeq X Plus platform using paired-end sequencing (2 × 150 bp).

### Quantification of differentially expressed genes

Raw paired-end mRNA-seq reads were assessed for quality using FastQC [47], trimmed using fastp (minimum length 30 bp, average quality ≥30) [48], and aligned to the ‘*Methylocystis sumavensis’* isolate genome (GCA_052186135.1) using HISAT2 [49]. Alignments were processed with SAMtools [50], and gene-level read counts were generated using featureCounts [51] based on coding sequence (CDS) annotations. Differential gene expression at pH 5.0 and pH 9.0 relative to the reference condition (pH 6.8) was assessed in DESeq2 [52] after filtering low-abundance genes (<10 total counts), using a negative binomial model with Benjamini–Hochberg correction. Genes with adjusted *p*-values (p_adj_) < 0.001 were considered significantly differentially expressed. Downstream visualisation and exploratory analyses were conducted in R using TPM and log₂(TPM + 1) values. Custom scripts developed for this study were informed by the transcriptomic analysis workflow established by Dreer et al. [53]. Functional categorisation was performed using Clusters of Orthologous Genes (COG) annotations. For each COG category, the numbers of differentially expressed genes that were up- or downregulated were compared for pH 5.0 vs pH 6.8 and pH 9.0 vs pH 6.8 using exact binomial tests against an expected 1:1 ratio and Fisher’s exact tests relative to all other COG categories; *p*-values were adjusted by the Benjamini–Hochberg method.

## Results and Discussion

### Cultivation, isolation, and cell morphology of a new Methylocystis species

For the enrichment of peatland methanotrophs, mineral medium was inoculated with pre-incubated top peat soil from a pristine minerotrophic fen in the Šumava Forest National Park, Czechia. Presence of methanotrophs was confirmed via PCR and CH_4_ oxidation was observed within serial dilutions after one week of incubation. Following plating and repeated streaking of single colonies, we obtained an active methanotrophic enrichment in liquid NMS medium. Since no growth of methanotroph taxa on solid medium could be achieved, the enrichment was subjected to random single-cell sorting into 96-well plates, which has successfully been applied to isolate other fastidious bacteria [54, 55]. Upon one month of incubation after sorting, growth was observed in multiple wells, and the cultures were identified as belonging to the genus *Methylocystis* with identical 16S rRNA genes. Absence of growth in organic medium and genome sequencing confirmed that the cultures were axenic.

Scanning electron microscopy revealed that the cells of the *Methylocystis* organism were predominantly single short rods measuring 1.4–1.7 µm in length and 0.5–0.7 µm in width with smooth surfaces similar to other *Methylocystis* species (Fig. 1A) [22, 56, 57]. High-resolution imaging (65,000×) showed no visible appendages (pili or flagella) or significant extracellular matrix, though such fragile structures may have been lost or damaged during dehydration or critical-point drying [58].

**Figure 1.**
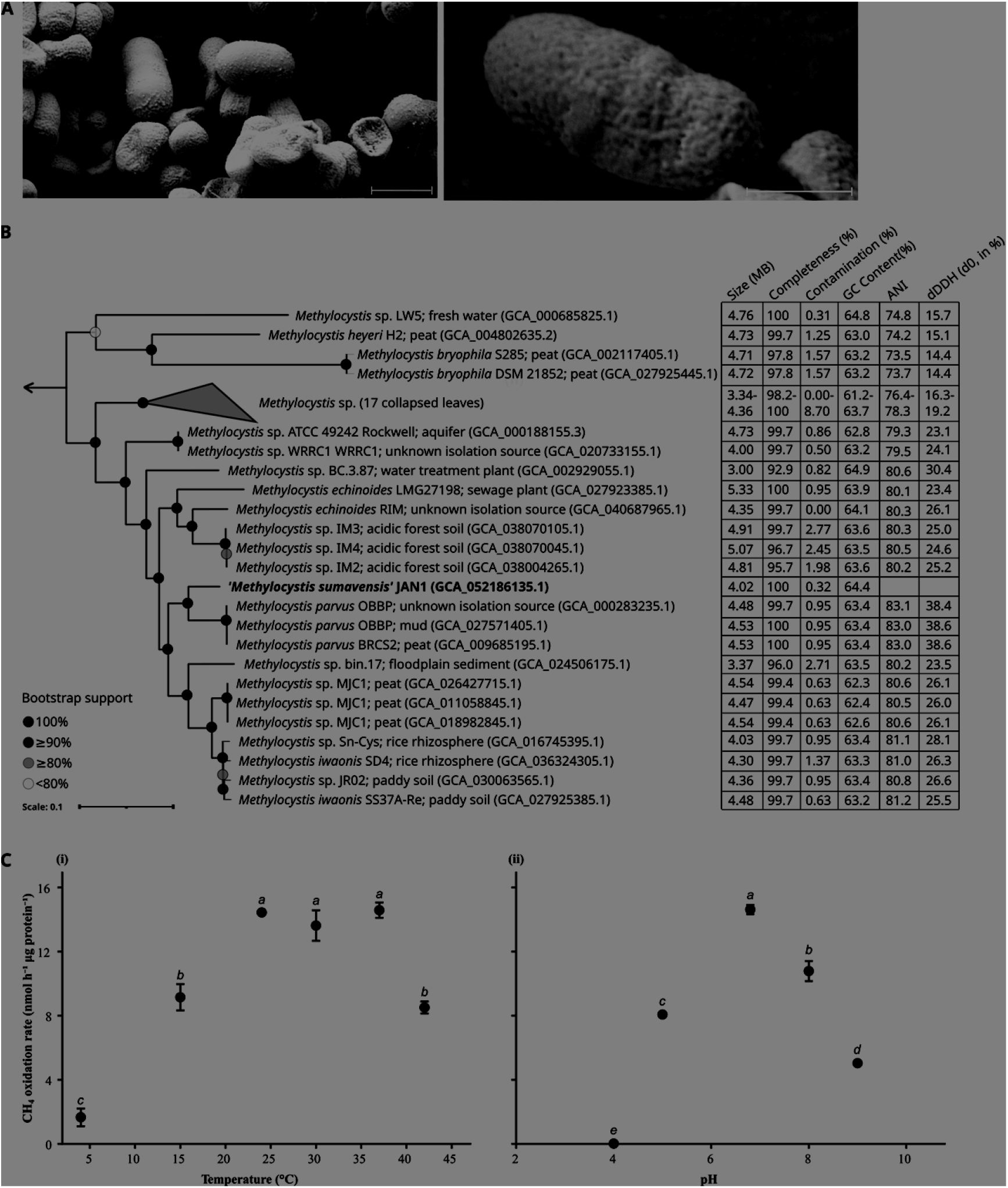
Morphology, phylogenomics, and physiology of ‘*Methylocystis sumavensis’*. **(A)** Scanning electron micrographs images of the newly cultured ‘*M. sumavensis’* showing short, rod-shaped cells at (i) low-magnification (18,231×) with 1 µm scale bar and (ii) higher resolution (65,000×) with 500 nm scale bar. **(B)** Phylogenomic placement and genomic characteristics of ‘*M. sumavensis’* and selected *Methylocystis* species. Names are followed by isolation source and the genome accession number in brackets. The maximum-likelihood phylogenomic tree shows the phylogenetic position of the newly cultured species (highlighted in bold) within the genus *Methylocystis*. The tree was constructed using the concatenated alignment of 71 single-copy core proteins with the LG+F+R3 substitution model and 1,000 ultrafast bootstrap replicates. Bootstrap support is indicated by coloured circles at nodes. The genome of *Methylomonas denitrificans* FJG1 (GCF_000785705.2) was used as an outgroup. **(C)** CH₄ oxidation rates by ‘*M. sumavensis’* tested at different temperatures (i) and pH (ii). Bars represent the average ± standard error of triplicate incubations. Different letters above bars indicate statistically significant differences (p < 0.05).

Long-read whole-genome sequencing of the newly cultured *Methylocystis* organism generated 166,564 reads. All reads were resolved into a single contiguous sequence, and *de novo* genome assembly resulted in only one complete circular chromosome with a length of 4.02 Mbp (GenBank assembly GCA_052186135.1) and a GC content of 64.4%. Additional genomic features are provided in Table S2. Based on GTDB taxonomy, the isolate was assigned to the genus *Methylocystis* but could not be assigned to a described species. Phylogenomic analysis placed the new *Methylocystis* isolate close to *Methylocystis parvus* clades but distinct from all currently described *M. parvus* strains (Fig. 2B) and phylogenetic analyses of the 16S rRNA gene (Fig. S1) and PmoA sequences showed a similar clustering pattern (Fig. S2). Genome-wide ANI (83.1%) and dDDH (38.4%) analyses against the closest relative, *Methylocystis parvus* OBBP (GCF_000283235.1), were below the species level thresholds (>95% ANI, >70% dDDH) [59], confirming the isolate as a novel species (Fig. 1; Tables S2–S3). We therefore propose the name ‘*Methylocystis sumavensis’* strain JAN1 after its place of isolation. The proposed name has been registered in the SeqCode Registry with GCA_052186135.1 as the nomenclatural type, and will be assigned to an official Register List upon validation.

**Figure 2.**
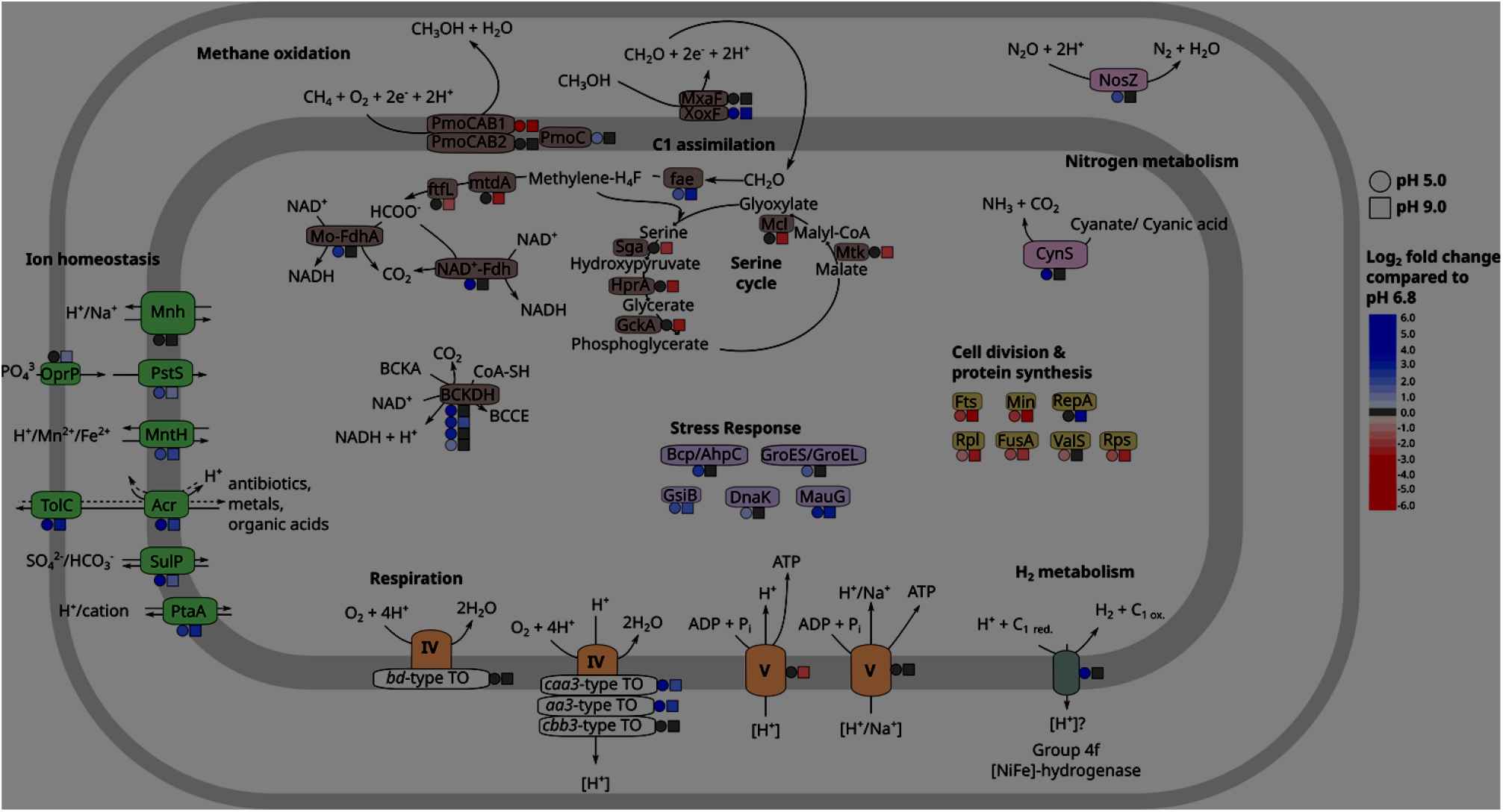
Schematic representation of the core metabolic and pH stress-response network of ‘*M. sumavensis’*. Shown are differential transcriptomic responses at pH 5.0 and pH 9.0 relative to the growth optimum (pH 6.8). The figure highlights major pathways, including CH₄ oxidation, C₁ assimilation, denitrification, hydrogen metabolism, respiration, ion homeostasis, and energy conservation mechanisms. Coloured symbols next to protein complexes indicate significant changes in transcription at pH 5.0 (circles) and pH 9.0 (squares). Colour intensity denotes log₂ fold change relative to pH 6.8, with blue indicating upregulation and red indicating downregulation. Roman numerals denote complexes of the electron transport chain. Abbreviations: PmoCAB1/2, particulate methane monooxygenase isoform 1 and 2; PmoC, orphan particulate methane monooxygenase subunit C paralog; MxaF/XoxF, Ca- and Ln-dependent methanol dehydrogenases; Fae, 5,6,7,8-tetrahydromethanopterin hydrolyase; MtdA, methylene-tetrahydromethanopterin dehydrogenase; FtfL, formate–tetrahydrofolate ligase; Mo-FdhA, molybdenum-dependent formate dehydrogenase catalytic subunit A; NAD⁺-Fdh, NAD⁺-dependent formate dehydrogenase (soluble formate dehydrogenase); Sga, serine–glyoxylate aminotransferase; HprA, hydroxypyruvate reductase; GckA, glycerate kinase; Mcl, malyl-CoA lyase; Mtk, malate-CoA ligase subunit alpha; BCKDH, branched-chain alpha-keto acid dehydrogenase complex; BCCE, branched-chain acyl-CoA esters; OprP, phosphate-selective porin; PstS, phosphate ABC transporter substrate-binding protein; MntH, Nramp-family divalent metal transporter (Mn²⁺/Fe²⁺); Mrp, multisubunit Na⁺/H⁺ antiporter complex; Acr, RND-family multidrug efflux transporter; TolC, outer membrane channel of the AcrAB–TolC efflux system; SulP, sulfate/bicarbonate transporter; PtaA, proton/cation antiporter; Bcp, bacterioferritin comigratory protein (peroxiredoxin); AhpC, alkyl hydroperoxide reductase subunit C; GroES, co-chaperonin GroES; GroEL, chaperonin GroEL; DnaK, molecular chaperone DnaK (Hsp70); GsiB, general stress protein GsiB; MauG, methylamine utilization protein MauG; Fts, cell division protein; Min, septum site-determining protein; RepA, replication initiation protein; Rpl, large-subunit ribosomal proteins; Rps, small-subunit ribosomal proteins; FusA, elongation factor G; ValS, valyl-tRNA synthetase; NosZ, nitrous oxide reductase; CynS, cyanase; Aa3-type TO, cytochrome c oxidase subunits of aa₃-type; Caa3-type TO, cytochrome c oxidase subunits of caa₃-type; Cbb3-type TO, cytochrome c oxidase subunits of cbb₃-type; Bd-type TO, cytochrome bd-type quinol oxidase subunits.

### ‘Methylocystis sumavensis’ is an acidotolerant mesophile

CH_4_ was actively consumed by ‘*M. sumavensis*’ at temperatures between 4 and 42°C, with optimal activity observed between 24–37 °C at a rate of 14.2 ± 0.51 nmol CH_4_ µg protein^-1^ · h^-1^ (Fig. 1C(i)). The observed oxidation activity at 42 °C was unexpected, particularly given the temperate origin of the isolate, as no other *Methylocystis* spp. have, to our knowledge, been reported to be active above 37 °C [22, 24, 56, 60–63], except for isolates from geothermal environments [64]. These findings show that the isolate is mesophilic and may be most active *in situ* during warmer periods.

When incubated across a range of pH conditions, ‘*M. sumavensis*’ showed CH_4_ oxidation activity across a broad pH range of 5.0–9.0 (Fig. 1C(ii)), with peak activity at pH 6.8 (14.6 ± 0.29 nmol CH_4_ µg protein^-1^ · h^-1^). This is consistent with the reported activity range of other *Methylocystis* spp. isolated from acidic habitats [22, 24]. Surprisingly, despite rapid CH_4_ consumption of the incubated peatland soil with a pH of 3.9–4.3 [18], no CH₄ oxidation by the axenic strain was detected at pH 4.0 under laboratory conditions (Fig. 1C(ii)). It remains to be verified whether ‘*M. sumavensis*’ can oxidise CH_4_ below pH 5 *in situ.* Possibly, in nature, members of the peat microbial community aid in pH stress alleviation [65], or under the tested laboratory conditions ‘*M. sumavensis*’ could not fully use its pH stress response capacity. Notably, in all incubations, including pH 5.0 and pH 9.0, the pH consistently shifted toward approximately 6.4 between pH adjustments, but not in the abiotic control (Fig. S3). This suggests active environmental pH modulation by the strain, possibly through metabolic buffering or ion exchange mechanisms.

### ‘Methylocystis sumavensis’ encodes a diverse repertoire for energy conservation and low pH tolerance’

The genome of ‘*M. sumavensis’* encodes a complete CH_4_ oxidation pathway, including both particulate methane monooxygenase (pMMO) isozymes: canonical pMMO1 (two *pmoCAB*1 clusters) and pMMO2 (*pmoCAB*2; Fig. 2, Table S4). Only half of the *Methylocystis* genomes analysed in this study possess both pMMO isozymes. This suggests a potential for atmospheric CH_4_ oxidation by ‘*M. sumavensis*’ [66], a hypothesis requiring experimental validation. Additionally, two singleton *pmoC* homologues, located outside the canonical structures, were identified in the ‘*M. sumavensis*’ genome (Fig. S4). Such orphan *pmoC* genes are typical in *Methylocystis* genomes, although their precise function is still unclear [67, 68]. No sMMO or pMMO-like (*pxmABC*) loci were detected in ‘*M. sumavensis*’ (Table S4), though these are respectively present in 18 and 24 of the 40 other *Methylocystis* genomes analysed, including those from acidic habitats.

Like most other *Methylocystis* spp., the genome of ‘*M. sumavensis*’ encodes the calcium-dependent methanol dehydrogenase (MxaF) as well as four copies of the lanthanide-dependent variant (XoxF) (Fig. 2, Table S4), both of which mediate methanol oxidation to formaldehyde [69, 70]. One of the *xoxF* homologues encodes a XoxF-2 type, whereas the other three encode the XoxF-5 type. Multiple *xoxF* homologues indicate functional diversification, regulatory differences, or substrate specialisation, with isoforms varying in kinetics, metal cofactor preference (La³⁺ vs. Ce³⁺), and responses to lanthanide availability [71–74].

Likewise typical for the genus, ‘*M. sumavensis’* possesses four different types of terminal oxidases. Specifically, these are three heme-copper cytochrome c oxidases of the aa_3_-, caa_3_-, and the high-affinity *cbb_3_*-type and a low-redox-potential high-affinity cytochrome *bd*-type ubiquinol oxidase (Fig. 2, Table S4). Together, they likely span a wide range of oxygen affinities, assumed to support respiration from fully oxic to micro-oxic conditions [75, 76].

Key genes for denitrification are present in ‘*M. sumavensis*’, including those encoding nitrite reductase (NirK, Table S4). Importantly, it possesses a complete *nos* gene cluster (*nosRZDFYX)* from a clade I nitrous oxide reductase (NOS) as defined by Hallin et al. [77] for the final denitrification step: reduction of nitrous oxide (N_2_O) to dinitrogen (N_2_). Similar *nos* gene clusters were found only in five other publicly available *Methylocystis* genomes: *M. echinoides* LMG27198, the species cluster *M.* sp. IM2, IM3, and IM4, and *M.* sp. SC2 (on a plasmid) ([78, 79]; Table S4). Although direct physiological assays are still pending, the presence of clade I *nos* genes suggests that ‘*M. sumavensis*’ may act as an N_2_O sink, reducing N_2_O to N_2_ in acidic peat environments, likely coupling this process to CH_4_ oxidation.

‘*M. sumavensis*’ has the potential to exploit energy sources other than CH_4_, as its genome encodes two [NiFe]-hydrogenases: a membrane-bound group 4f (Ech) and a cytosolic group 3b (Frh), along with maturation and regulatory proteins (Fig. 2, Table S4, Fig. S5, Fig. S6). The group 4f complex, though uncharacterised, is predicted to couple formate oxidation to H_2_ production and proton translocation under anoxic conditions [80]. HydDB-based identification and classification of catalytic subunits across the 40-genome dataset showed that group 4f hydrogenases are widespread in *Methylocystis*, occurring in 38 of 40 publicly available genomes. To our knowledge, this is the first report of a conserved group 4f [NiFe]-hydrogenase in *Methylocystis*, as a recent comparative study reported only groups 1a, 1d, 1h/5, 1i, 2b, 2c, 3b, and 3d in the genus [81]. Moreover, alignment of 39 *Methylocystis* group 4f catalytic subunits, including that of our isolate, with 23 HydDB group 4f reference sequences revealed a highly conserved *Methylocystis*-specific sequence motif (e.g. PLHQIPVGP…ANGExVVRL…AARVS) with multiple amino acid positions differing consistently from the non-*Methylocystis* group 4f consensus (Figure S7). The catalytic subunit of the group 3b hydrogenase, which likely mediates NADPH-dependent reversible H_2_ evolution, clusters with homologues from other *Methylocystis* and *Methylosinus* species (Fig. S5 and Fig. S6).

Adjacent to the Frh loci, we identified a 20-kDa electron-transfer subunit and a small Fe-S protein previously annotated as anaerobic sulphite reductase subunit B (AsrB). Given its genomic context with FrhA and FrhG, we interpret it as the electron-transfer subunit of the [NiFe]-hydrogenase, consistent with previous suggestions of sulfhydrogenase-like complex structures [82].

Acidophilic and acidotolerant microorganisms need to maintain near-neutral intracellular pH to ensure optimal protein function despite low external pH, which results in a high transmembrane pH gradient (ΔpH; much lower outside than inside the cell) and a reversed transmembrane electrical potential (Δψ; positive inside relative to the outside of the cell) [83, 84]. Whereas the former is beneficial for energy conservation (proton motive force (PMF) increases with ΔpH), the reversed Δψ poses a significant bioenergetic challenge. We identified several genomic features that may aid ‘*M. sumavensis’* in maintaining pH homeostasis via proton extrusion, such as a multisubunit H⁺/Na⁺ antiporter (MrpA–G) that exports protons in exchange for Na⁺ (Fig. 2, Table S6). This complex, which was found in only 7 of 41 analysed *Methylocystis* genomes, may act in concert with other antiporters (ClcA, ChaAB, NhaP), the membrane-bound pyrophosphatase OVP1 and systems involved in membrane integrity and stress tolerance (Table S5 and S6).

Additionally, ‘*M. sumavensis*’ encodes both canonical H⁺- and predicted H⁺/Na⁺-translocating ATP synthases (Fig. 2, Table S4, Fig. S8–S9). This H⁺/Na⁺-translocating variant is likewise present in a few other *Methylocystis* spp., particularly those isolated from acidic habitats, and has high sequence similarity and predicted 3D structure with characterised H⁺/Na⁺ -translocating AtpE subunits from bacteria shown to perform Na^+^-dependent ATP synthesis *[85, 86]* (Fig. S9, Table S7A and 7B). This dual ATP synthase system suggests metabolic flexibility in energy conservation, allowing ATP generation via chemo-osmotic coupling under proton and, potentially, sodium ion gradients [86, 87], which may be advantageous under the variable pH and pore water chemistry characteristic of minerotrophic peatland habitats [88, 89].

### Transcriptomic responses to pH stress

To link genomic potential with physiology and identify metabolic mechanisms underlying pH stress tolerance, we compared gene expression of ‘*M. sumavensis’* under acidic (pH 5.0) and alkaline (pH 9.0) conditions relative to the standard cultivation condition (pH 6.8). Although alkaline conditions are not representative of the native peatland environment, pH 9.0 was included as a physiological boundary condition to contrast responses to proton excess. We identified 519 significantly up- and 200 downregulated genes at pH 5.0, and 445 significantly up- and 457 downregulated genes at pH 9.0 (Fig. 3; Tables S8, S9).

**Figure 3.**
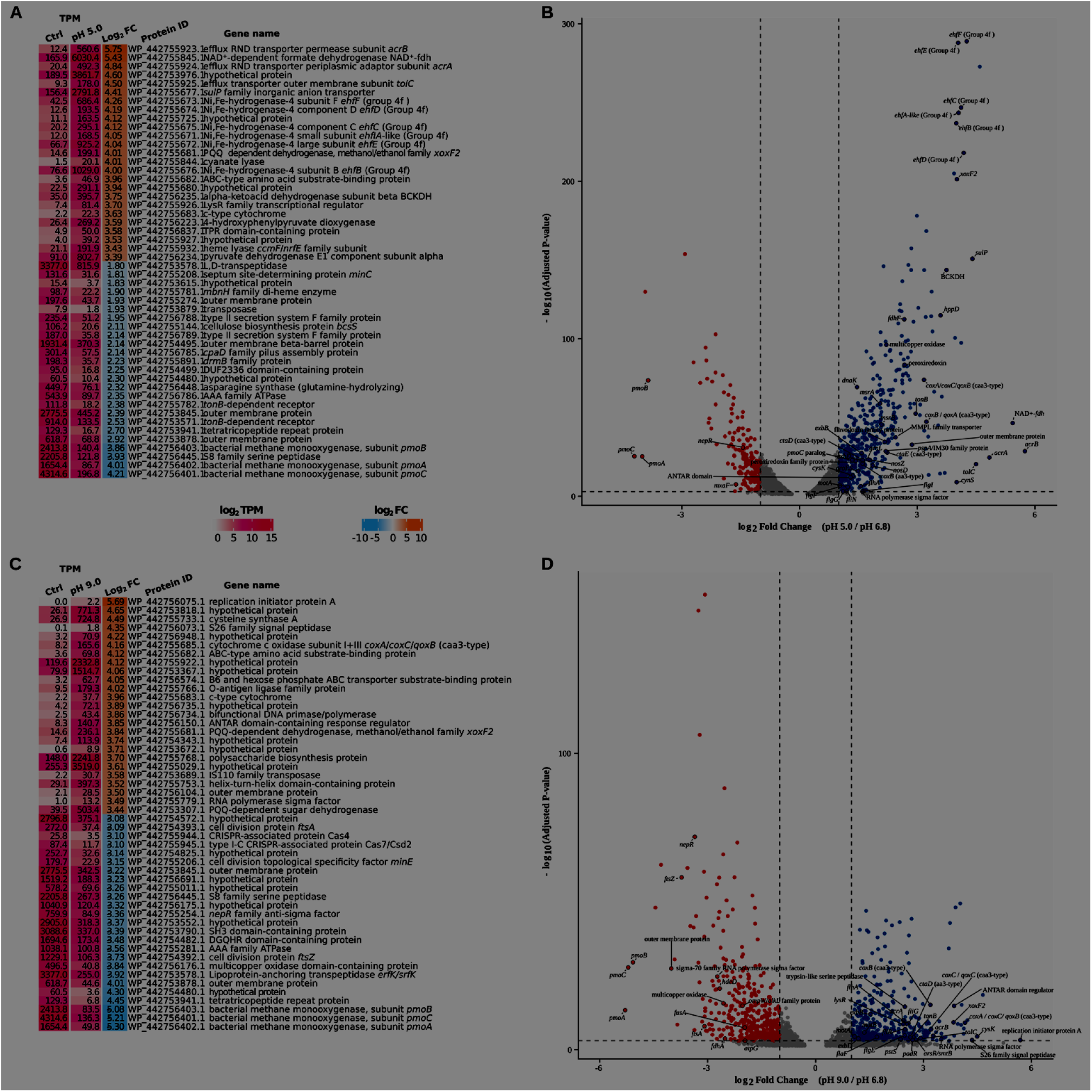
Differentially expressed genes in ‘*M. sumavensis*’ at pH 5.0 and pH 9.0 relative to the growth optimum (pH 6.8). Panels (A) and (C) show the top 50 up- and downregulated genes at pH 5.0 and pH 9.0, respectively, relative to the reference condition pH 6.8, ordered by log₂ fold change (log₂FC), and expression levels (transcripts per million, TPM) for both control and treatment conditions. TPM values are colour-coded based on log₂-transformed expression levels. Panels (B) and (D) show the corresponding volcano plots for pH 5.0 and pH 9.0, respectively. In the volcano plots, dashed vertical lines mark log_2_ fold change thresholds of -1 and 1, and the dashed horizontal line marks the significance cutoff of adjusted P-value < 0.001. Red dots = down-regulated; blue dots = up-regulated; grey dots = not significant. Selected genes are labelled for emphasis. Abbreviations: AcrA/AcrB/TolC efflux system; BCKDH, branched-chain alpha-ketoacid dehydrogenase complex; Ehf, group 4f [NiFe]-hydrogenase; Fdh/FdhF, formate dehydrogenase; MxaF, methanol dehydrogenase [cytochrome c] subunit; NosZ/NosD, nitrous-oxide reductase and maturation protein; PmoA/PmoB/PmoC, particulate methane monooxygenase subunits; PspA, PspA/IM30 family protein; MMPL, MMPL family transporter; MsrA/MsrB, methionine sulfoxide reductases; GroL/GroES/DnaK, chaperones; Flg/Flh/Fli/Mot/Exb/TonB, flagellar and TonB-system proteins; Cta/Cox, cytochrome c oxidase subunits; PstS, phosphate ABC transporter substrate-binding protein; AtpG, ATP synthase gamma subunit; FtsA/FtsZ, cell division proteins. For the full dataset, see Tables S8 and S9.

Functional categorisation of expressed genes using the COG classification revealed contrasting strategies across pH conditions (Fig. S10; Tables S8 and S9). At pH 5.0, genes assigned to several COG categories, including energy production and conversion, amino acid transport and metabolism, lipid transport and metabolism, inorganic ion transport and metabolism, and secondary metabolite biosynthesis, were significantly biased towards upregulation (Fig. S10A, Table S10). This pattern is consistent with increased energetic and cellular maintenance demands under acidic conditions [26]. In contrast, at pH 9.0, genes associated with cell cycle control, cell division, and chromosome partitioning, translation, ribosomal structure and biogenesis, and cell wall/membrane/envelope biogenesis were significantly biased toward downregulation (Fig. S10B, Table S10). This pattern suggests a stress-acclimation response characterised by suppression of growth-related processes and a shift toward basic maintenance and cell envelope homeostasis under alkaline conditions.

In both conditions, a substantial fraction of the differentially upregulated genes belonged to the unknown-function category (COG S), suggesting that currently uncharacterised mechanisms may contribute to pH stress adaptation in ‘*M. sumavensis*’.

### Diverse transcriptional responses coordinate acid stress (pH 5.0) tolerance

We observed substantial downregulation of pMMO1 genes (*pmoCAB1*) at pH 5.0 (Fig. 2, Fig. 3A and 3B; Table S8; log_2_FC = -3.9 to -4.5). pMMO1 was the most highly transcribed MMO isozyme and likely acted as the primary MMO under all tested pH conditions. Its reduced transcription under acid stress is consistent with the observed lower CH_4_ oxidation under the these conditions. However, we detected modest induction of an orphan *pmoC* homolog (log_2_FC = +1.3), whose function remains unknown. We further observed a shift in methanol dehydrogenase transcription patterns with a strong upregulation of one of the lanthanide-dependent methanol dehydrogenases (XoxF2-type; log_2_FC = +4.0) at pH 5.0 versus downregulation of the Ca-dependent type (MxaF; log_2_FC = -1.65), which was the primary methanol dehydrogenase at neutral pH. This can be explained by the reported higher rate and substrate affinity of certain XoxF-methanol dehydrogenase and their ability to directly convert methanol to formate without the need for methanopterin- or folate-dependent enzymes [74], which may reduce the biosynthetic burden relative to the Ca-dependent MxaF system. Together, these results demonstrate differential regulation of proteins involved in CH_4_ and methanol oxidation during acid stress.

Among the most strongly upregulated genes under the acid stress condition were all subunit genes encoding a canonical tripartite RND-family efflux system (AcrAB–TolC-like; log_2_FC = +4.6–5.8; Fig. 2, Fig. 3A and 3B; Table S8). This PMF–driven efflux complex is characterised by broad substrate poly-specificity and the capacity to export structurally diverse toxic compounds and metabolic by-products [90]. Comparative screening of 40 *Methylocystis* genomes revealed that AcrAB–TolC is not universally conserved in the genus but occurs more frequently in acid-associated lineages: merely 15 genomes (36.6%) encode a complete system, and 11 of these (∼73%) originate from peat or acidic forest soils (Table S11). Several additional RND permeases and adaptor proteins from other genomic loci were also moderately upregulated (Table S8), indicating activation of multiple efflux systems. Even though the specific compounds exported under these conditions remain unknown, this finding suggests that efflux forms part of the pH stress response in ‘*M. sumavensis’*, consistent with observations in other bacteria where RND-type efflux pumps were induced under acidic stress conditions [91].

Low pH further induced broad upregulation of respiratory and energy metabolism genes in ‘*M. sumavensis’*, which indicates increased electron flow (Fig. 2, Fig. 3A and 3B; Table S8). Notably, these included the complete membrane-bound group 4f [NiFe]-hydrogenase complex (*ehfA-F*; log_2_FC = +4.2–4.5), two formate dehydrogenases (log_2_FC = +2.67 – 5.43), all three subunits of the branched-chain α-keto-acid dehydrogenase complex (BCKDH; log_2_FC = +3.8), and different terminal oxidase subunits, including the caa_3_-type (log_2_FC = +2.2–3.2) and aa_3_-type (log_2_FC = +1– 1.6). BCKDH contributes to branched-chain fatty acid synthesis involved in maintaining membrane functionality [92]. Moreover, upregulation of BCKDH and formate dehydrogenases is expected to increase the supply of reducing equivalents (*e.g.*, NADH), thereby sustaining respiratory flux and supporting PMF and ATP synthesis. Concomitant induction of multiple terminal oxidases further suggests reinforcement of electron transport capacity, indicating a coordinated upregulation of respiratory metabolism, most likely associated with maintenance of cytoplasmic pH homeostasis under acid stress. Similarly, modulation of respiratory flux and proton-translocating complexes has been associated with cytoplasmic pH homeostasis and bioenergetic balance under pH stress in other bacteria [28, 93–95,].

The induction of the group 4f [NiFe]-hydrogenase (Ech-type), which forms a membrane-associated complex evolutionarily related to respiratory complex I and is thought to couple proton reduction to ion translocation [96], suggests a link between hydrogen metabolism and chemiosmotic energy conservation and redox balancing in ‘*M. sumavensis’* at acidic pH. A highly similar mechanism has been observed for *E. coli*, which requires its related group 4a [NiFe]-hydrogenase for survival at acidic pH, albeit during anaerobic growth [97]. However, we could not obtain evidence for H_2_ evolution under the oxic incubation conditions applied here (Table S12), which does not support activity of the hydrogenase in fully oxic conditions despite significant upregulation of the hydrogenase genes. The detection limit of the method used was 40 ppmv H_2_, thus, if H_2_ was produced, it was below this reliable quantification range. Future experiments examining anoxic conditions and the supply of external C1 compounds are needed to verify whether the 4f hydrogenase plays a role in the activity of ‘*Methylocystis sumavensis’* at low pH.

Finally, we detected increased transcription of oxidative stress defence, membrane stabilisation, proteostasis systems, and auxiliary redox-active proteins in response to acid stress (Supplementary Text). Together, these varied transcriptional changes indicate a coordinated acid stress response that couples respiratory chain restructuring with redox balancing, cellular maintenance, and membrane reinforcement to sustain energy conservation at low pH.

### Transcriptional response to alkaline stress (pH 9.0) largely reflect growth suppression, damage repair, and ion homeostasis

Overall, the transcriptional changes observed at pH 9.0 suggest a regulatory response that prioritises ionic balance, envelope integrity, and genome stability at the expense of methanotrophic growth (Fig. 2, Fig. 3C and 3D, Fig. S9, Table S9). Although aa3- and caa3-type cytochrome c oxidase subunits were induced (log_2_FC = +2.2–4.2), suggesting a concerted effort to maintain terminal electron transfer capacity and the PMF under alkaline stress, transcripts encoding growth-related functions were among the most repressed. This included transcripts for multiple cell division proteins including FtsA and FtsZ, ribosomal proteins (*rpl*, *rps* genes), translation factors (*fusA*), ATP synthase subunits (*atpE*, *atpG*), the primary CH_4_-oxidation complex, pMMO1 (log_2_FC ≈ −5.1 to −5.3), and several core serine-cycle genes, including *glyA*, *ppc*, *pckA*, and a putative gckA-like glycerate kinase (Fig. 2, Fig. 3C and 3D, Table S9). Downregulation of these genes highlights a metabolic shift away from energy-intensive biosynthesis toward a maintenance-oriented physiological state. Interestingly, similar to the pH 5 condition, the transcription of the same lanthanide-dependent methanol dehydrogenase, XoxF2, was highly increased (log_2_FC = +3.84), suggesting its preferential utilisation under both acidic and alkaline pH stress.

Further reflecting the physiological stress experienced by the cells, three different genes coding for NepR family anti-sigma factors were highly downregulated (log_2_FC = -3.4 to -1.7). These proteins act as negative regulators that bind to and inactivate sigma factors responsible for global transcriptional control of gene expression under specific stress conditions [98]. The reduced transcription of anti-sigma factors indicates the activation of a broader stress response. In contrast, general regulatory and genome maintenance functions were strongly upregulated, indicating a shift towards cellular preservation. Key induced genes included an RNA polymerase sigma factor, *rpoE* (log_2_FC = +3.5), an ANTAR-domain response regulator (log_2_FC = +3.9), and genes involved in DNA repair (DNA ligase D, Ku protein, ATP-dependent DNA ligase, Y-family DNA polymerase, DNA polymerase IV dinB, RecT recombinase; log_2_FC = +1.2–2.9). These responses are consistent with a broader stress-adapted state where transcriptional control and genome integrity are prioritised over cell division [27]. In this context, the finding that the gene for the replication initiator protein A (*dnaA*) was the most strongly upregulated gene at pH 9 (log_2_FC = +5.7) seems contradictory. We speculate that, rather than indicating active replication, it may be related to stress-associated replication control or DNA maintenance processes under alkaline conditions, although its precise role remains unclear.

Under alkaline conditions, active inward transport of protons and accumulation of compatible solutes that support cellular homeostasis under alkaline conditions are crucial adaptations and usually involve activation and transcriptional upregulation of various transporters [27]. Consistently, we observed a strong upregulation of genes for several multi-subunit cation and nutrient transporters (Fig. 2, Fig. 3C and 3D, Table S9), including the tripartite RND-family efflux system (AcrAB– TolC-like; log_2_FC = +2.4 to +3.3) which was also activated at pH 5.0, as well as additional RND permease and outer membrane efflux components (log_2_FC = +2.5 to +3.1). Other upregulated genes included subunits of an ABC glycine/betaine transporter (log_2_FC = +2.2 and +2.1), a P-type ATPase (log_2_FC = +2.9), and a Nramp-family proton/divalent metal symporter (log_2_FC = +2.14). Surprisingly, none of the genes encoding the multi-subunit H⁺/Na⁺ antiporter MrpA–G, which is considered to be critical at elevated pH in alkaliphilic bacteria [28, 99], were among the significantly upregulated genes. Possibly, the sodium levels in the medium were not sufficient to support this antiporter and/or the regulation of this system is not linked to elevated pH alone in ‘*M. sumavensis’* [100]. Additionally, we observed strong upregulation of genes involved in generating acids and antioxidant/ROS detoxifying systems, flagellar synthesis, and cell envelope modification (Supplementary Text).

## Conclusion

Even though the level of pH stress was not the same at pH 5.0 and 9.0, as reflected by the higher CH_4_ oxidation activity maintained at pH 5.0 (Fig. 1), our results collectively suggest that acid stress in ‘*M. sumavensis’* elicits an energetically demanding, redox-active compensatory response, whereas alkaline stress promotes a growth-suppressive regulatory strategy focused on cellular maintenance. Nevertheless, several adaptive responses, including cell-envelope remodelling and redox balancing, were shared between both conditions, although they were mediated through distinct mechanisms at low and high pH.

### Taxonomic consideration of ‘*Methylocystis sumavensis*’

*Methylocystis sumavensis* (su.ma.ven’sis L. fem. adj. sumavensis, of Šumava, the region in the Czech Republic from which the fen peatland samples containing this organism were obtained). An aerobic CH_4_-oxidising rod-shaped bacterium isolated from the surface layer (1–10 cm) of a fen peatland located in the Šumava Forest National Park, Czechia. Phylogenetically affiliated with the family *Methylocystaceae*, phylum *Proteobacteria*. Cells are ∼1.5 μm long Gram-negative rods. The genome consists of a single chromosome of 4,022,259 bp. The DNA G + C content is 64 mol%. The strain was routinely cultured in liquid minimal mineral medium with nitrate as a N source and 10% CH_4_ as energy and C source. Although isolated in pure culture, strain JAN1 cannot currently be deposited in recognised culture collections due to preservation challenges, making formal validation under International Code of Nomenclature of Prokaryotes criteria pending.

The genome of ‘*Methylocystis sumavensis’* JAN1, available under GenBank assembly GCA_052186135.1, is the designated nomenclatural type for the species. ANI comparisons with closely related *Methylocystis* genomes are below 95%, supporting the delineation of this isolate as distinct from other species in the genus.

## Acknowledgments

The authors thank Tomáš Picek and Zuzana Urbanová for access to the field site. We further acknowledge the BC CAS core facility LEM supported by MEYS CR (LM2023050 Czech-BioImaging and OP VVV CZ.02.1.01/0.0/0.0/18_046/0016045), for granting us access to instrumentation and support with sample preparation and imaging. We thank Ondřej Honc (Biotechnology and Biomedicine Center of the Academy of Sciences and Charles University in Vestec, BIOCEV) for assistance with cell sorting using FACS. Finally, we thank Travis Meador (Biology Centre CAS) for molecular hydrogen quantification. J.A.N., G.I., M.W., and A.D. were supported by a JuniorStar grant from the Czech Science Foundation (CSF) (grant number 21-17322 M) awarded to A.D.

## Data availability

The genome sequence of ‘*M. sumavensis’* JAN1 has been deposited under BioProject ID PRJNA1216969 with accession number GCA_052186135.1. The name ‘*Methylocystis sumavensis*’ has been registered in the SeqCode Registry (Register List – pending assignment), with genome GCA_052186135.1 designated as the nomenclatural type. Raw transcriptomic (mRNA-seq) data have been deposited in the NCBI Sequence Read Archive under the same BioProject. All downstream analysis scripts, processed expression matrices, differential expression output tables, and annotation files (including DRAM and COG outputs) are available in a public GitHub repository: https://github.com/Justan6/JAN1_draft_genome_metadata_and_analysis-scripts. DRAM-annotated protein sequences from all *Methylocystis* genomes and selected MAGs used in comparative analyses are available at https://zenodo.org/records/16927688

## Author contributions

J.A.N (conceptualisation, data curation, formal analysis, investigation, visualisation, writing – original draft, review & editing), M. W. (conceptualisation, investigation, enrichment establishment, writing – review & editing), G. I. (investigation, enrichment establishment), and A.D (conceptualisation, data curation, formal analysis, funding acquisition, writing – original draft, review & editing).

