## Supplementary figures and images for "Genomic, physiological, and transcriptomic mechanisms of pH stress adaptation and response in *Methylocystis sumavensis*, a novel peatland methanotroph"

### Supplementary text and figures

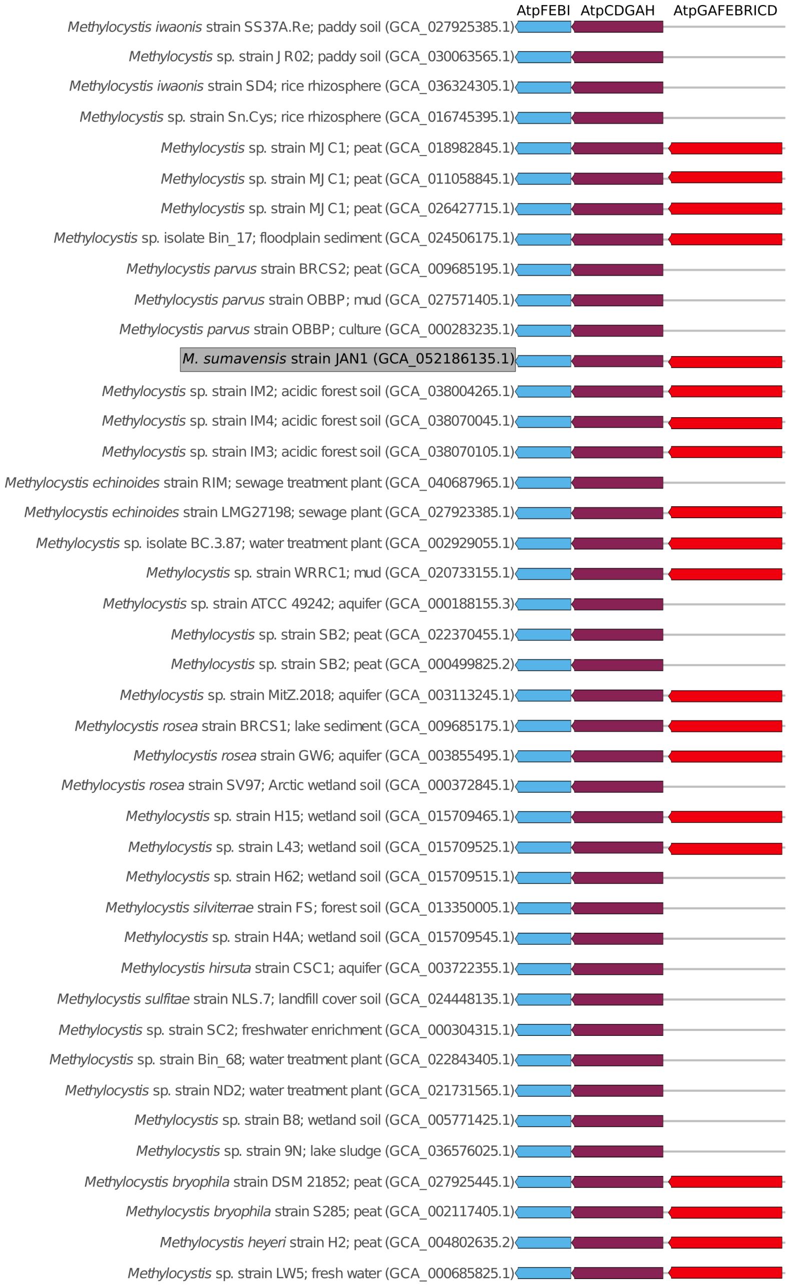
